# Ketosis Elevates Antioxidants and Enhances Neural Function Through Improved Bioenergetics: A ^1^H MR Spectroscopy Study

**DOI:** 10.1101/2024.10.22.619722

**Authors:** Helena van Nieuwenhuizen, Botond B. Antal, Antoine Hone-Blanchet, Andrew Lithen, Liam McMahon, Sofia Nikolaidou, Zeming Kuang, Kieran Clarke, Bruce G. Jenkins, Douglas L. Rothman, Lilianne R. Mujica-Parodi, Eva-Maria Ratai

## Abstract

Ketosis is known to alter the balance of neuroactive amino acids and enhance neural function when compared to a glycolytic condition. However, its influence on other metabolites, such as antioxidants and neural energy markers, and the mechanisms by which ketosis improves neural function remain unclear. Here, we measure the neurochemical effects of acute ketosis on the human brain using ultra-high-field ^1^H MR Spectroscopy (MRS) and investigate the subsequent impact on neural function through resting-state functional magnetic resonance imaging (rsfMRI). In a within-subjects design, N = 63 healthy adults from across the lifespan underwent ^1^H MRS and rsfMRI scans before and after consuming individually weight-dosed and calorically-matched ketone monoester or glucose drinks. Ketone monoester administration, but not glucose, significantly elevated cerebral antioxidants and energy markers while decreasing GABA, glutamate, and glutamine levels in the posterior cingulate cortex (PCC). Notably, increased bioenergetics, specifically an increase in total creatine, correlated with greater improvements in neural function as measured using rsfMRI. Our results integrate metabolic and functional neuroimaging findings, offering a comprehensive understanding of ketosis-induced changes in brain chemistry and functional network dynamics, yielding valuable insights into potential mechanisms by which ketosis imparts its neural benefits.

## Introduction

Ketone bodies, such as D-*β*-hydroxybutyrate (D-*β*HB), are naturally produced by the liver during periods of low carbohydrate availability, and function as an alternative fuel source to cells in the absence of glucose. Ketosis, whether achieved through a ketogenic diet or via exogenous ketone consumption, offers numerous benefits to brain health across a range of diseases and disorders. One of the most well-documented successes of the ketogenic diet is its ability to reduce the frequency and severity of seizures in children with drug-resistant epilepsy [12, 38]. Ketone bodies are also able to be utilized by insulin-resistant neurons, which may reverse and prevent brain aging and neurodegeneration caused by a lack of available neural energy in diseases such as diabetes [1, 8]. Additionally, a recent pilot study found an improvement in psychiatric symptoms following a ketogenic diet intervention in individuals with schizophrenia or bipolar disorder with existing metabolic abnormalities [51]. Previous work measuring metabolite concentrations in the brain under two metabolic conditions (ketotic and glycolytic) using ^1^H magnetic resonance spectroscopy (MRS) found acute oral administration of exogenous ketone monoester (d-*β*-hydroxybutyrate, or D-*β*HB) significantly reduced levels of both *γ*-Aminobutyric Acid (GABA) and glutamate (Glu) in the anterior and posterior cingulate cortices (ACC and PCC) of healthy adults. This effect was only observed following D-*β*HB administration and not following glucose administration, suggesting a unique impact of ketones on brain metabolism, particularly in older adults who showed a greater magnitude of these effects [20]. Finally, our previous research using resting-state fMRI has shown that the stability of functional communication across brain regions over time (*brain network stability* ) improves in response to both ketosis achieved through longer-term dietary conditions as well as acute oral administration of individually weight-dosed D-*β*HB [37].

Despite the observed benefits of ketosis on brain function, the precise impact of ketosis on cerebral antioxidants and energy markers, and how these changes potentially contribute to improved neurological outcomes and brain function, remain unknown. As both oxidative stress and mitochondrial deficits are associated with aging [25], neuroinflammatory illnesses [36], and psychiatric disorders [39, 45], understanding the effects of and mechanisms by which ketosis impacts these systems is important for understanding how it can mitigate the progression and severity of neurological disorders.

To this end, we conduct a more thorough analysis by increasing the sample size of our previous study from N = 26 to N = 63 participants, still sampled across the lifespan and within the PCC, allowing for a more robust examination of the neurochemical effects of ketone administration, especially for those metabolites that are naturally low in concentration in the brain. The posterior cingulate cortex (PCC), a key node in the default mode network, was selected as the continued region of interest for our ^1^H MRS measurements due to its high baseline metabolic rate [44] and susceptibility to metabolic changes associated with neurodegenerative disease, as evidenced by marked metabolic reduction in the PCC in individuals with very early Alzheimer’s disease [34]. Our expanded study not only replicates our original findings of decreased GABA and glutamate levels, but also finds increases in cerebral antioxidants and energy markers and examines the subsequent impact on brain function using resting-state functional magnetic resonance imaging (rsfMRI). These novel results provide valuable insights into the underlying mechanisms through which ketosis may improve the prognosis of neurological disorders, highlighting potential directions for future research and the development of potential therapeutic strategies.

## Methods

### Bolus Study

#### Study Population

^1^H MRS and resting-state fMRI (rsfMRI) data were collected from a cohort of healthy adults aged between 22 and 79 years. Detailed clinical and demographic characteristics for the study population can be found in Table 1. Exclusion criteria included MR contraindications for ultra-high-field imaging, diagnoses of psychiatric and/or neurological disorders, diagnoses of diabetes mellitus, insulin resistance (as measured via HOMA-IR/OGTT), history of brain injury, recreational drug use, severe alcohol use, use of medications that affect glucose and/or insulin utilization, and current or recent adherence to a low-carbohydrate or ketogenic diet. The study was registered as a clinical trial on ClinicalTrials.gov (identifier NCT04106882) and approved by the institutional review boards of Massachusetts General Hospital (2015P000652) and Stony Brook University (IRB2019-00208). Participants were recruited from the Boston metropolitan area through online and paper advertisements. All participants provided written informed consent.

**Table 1.**
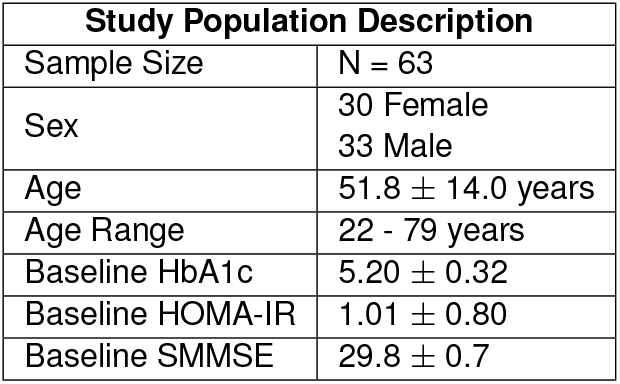
Clinical and demographic information of study participants. Includes clinical marker results collected at screening visit. Values given in mean *±* standard deviation.

| Study Population Description |  |
| --- | --- |
| Sample Size | N = 63 |
| Sex | 30 Female<br>33 Male |
| Age | 51.8 ± 14.0 years |
| Age Range | 22 - 79 years |
| Baseline HbA1c | 5.20 ± 0.32 |
| Baseline HOMA-IR | 1.01 ± 0.80 |
| Baseline SMMSE | 29.8 ± 0.7 |

**Table 2.** Descriptive statistics of metabolite concentrations pre- and post-bolus. A *∗* indicates a significant (p *<* 0.05) change in the metabolite concentration following the ketone bolus, while a *†* indicates a significant change in the metabolite concetration following the glucose bolus.

| Metabolite | Metabolite Concentrations Pre- vs. Post-Bolus |  |  |  |  |  |
| --- | --- | --- | --- | --- | --- | --- |
| | D-βHB Condition | | | Glc Condition | | |
|  | N | z | p-value | N | z | p-value |
| Asc* | 34 | 4.584 | 4.571E-06 | 30 | 1.329 | 0.184 |
| Asp | 56 | -0.865 | 0.387 | 47 | -0.488 | 0.626 |
| Cr* | 56 | -2.945 | 0.003 | 53 | -0.084 | 0.933 |
| Cr+PCr* | 57 | 2.769 | 0.006 | 53 | -0.261 | 0.794 |
| D-βHB* | 39 | 6.951 | 3.638E-12 | 30 | -1.018 | 0.309 |
| GABA* | 40 | -6.030 | 1.644E-09 | 50 | -0.239 | 0.811 |
| GABA/Glu* | 40 | -5.493 | 3.949E-08 | 38 | 1.935 | 0.053 |
| Glc<sup>†</sup> | 28 | 1.346 | 0.178 | 25 | 3.883 | 1.030E-04 |
| Glc+Tau<sup>†</sup> | 55 | 0.084 | 0.933 | 52 | 5.145 | 2.669E-07 |
| Gln* | 57 | -3.277 | 0.001 | 53 | -1.439 | 0.150 |
| Gln/Glu* | 57 | 3.524 | 4.256E-04 | 53 | -1.182 | 0.237 |
| Glu* | 57 | -6.050 | 1.446E-09 | 53 | -0.252 | 0.801 |
| Glu/GABA* | 40 | 5.135 | 2.823E-07 | 38 | -1.681 | 0.093 |
| Glu/Gln* | 57 | -3.492 | 4.796E-04 | 53 | 1.403 | 0.161 |
| Glu+Gln* | 57 | -5.732 | 9.898E-09 | 53 | -0.482 | 0.629 |
| GPC* | 57 | 6.503 | 7.865E-11 | 53 | 0.881 | 0.378 |
| GSH* | 57 | 3.341 | 8.349E-04 | 53 | -0.527 | 0.598 |
| Ins | 57 | 1.855 | 0.064 | 53 | -1.058 | 0.29 |
| Lac | 43 | -1.745 | 0.081 | 44 | -0.098 | 0.922 |
| Mac* | 57 | 2.316 | 0.021 | 53 | 0.04 | 0.968 |
| NAA | 57 | 0.107 | 0.915 | 53 | -0.013 | 0.989 |
| NAAG | 55 | -1.894 | 0.058 | 51 | -1.903 | 0.057 |
| NAA+NAAG | 57 | -0.497 | 0.619 | 53 | -0.350 | 0.727 |
| PCho | 4 | -1.534 | 0.125 | 34 | -0.236 | 0.813 |
| PCho+GPC | 57 | 0.814 | 0.415 | 53 | -0.695 | 0.487 |
| PCr* | 57 | 3.818 | 1.347E-04 | 53 | -0.421 | 0.674 |
| PE | 54 | -1.442 | 0.149 | 53 | -0.394 | 0.694 |
| sIns | 42 | -0.458 | 0.647 | 41 | -0.423 | 0.672 |
| Tau*<sup>†</sup> | 56 | -3.630 | 2.835E-04 | 53 | 2.226 | 0.026 |

#### Experimental Design

To investigate the impact of acute administration of D-*β*HB ketone monoester and calorically matched glucose on metabolite concentrations and brain network stabilities, resting-state fMRI and ^1^H MRS were collected from a cohort of healthy adults aged between 22 and 79 years who were tested under both a ketotic (ketone burning) and glycolytic (glucose burning) condition in a within-subjects experimental design. At an initial screening visit, participants first underwent a physical examination, an oral glucose tolerance test (OGTT), a metabolic blood panel, and a Standardized Mini-Mental State Examination (SMMSE) to confirm cognitive normalcy and healthy control status. Within 30 days of the initial screening visit, participants were tested twice (1-14 days apart), both times following an overnight fast (no food for at least eight hours before testing but unrestricted water). Following a baseline resting-state scan, participants consumed one of two fuel sources. During the ketotic session, they drank a ketone drink prepared using pure (R)-3-hydroxybutyl-(R)-3-hydroxybutyrate monoester (ΔG ketone ester, HVMN Inc., Miami, FL, USA) dosed at 395 mg/kg and diluted with water (volume ratio 1:1.6). During the glycolytic session, the same participants ingested a glucose bolus (Glucose Tolerance Test Beverages, Fisher Scientific, Inc.; Hampton NH) calorie-matched to the ketone monoester bolus. The order of the bolus was randomized. The resting-state scans were repeated 30 minutes after administration of the bolus, based on prior ^1^H MRS experiments indicating that peak glucose and ketone concentrations in the brain occur approximately 30 minutes post-consumption [37]. During the resting-state scans, a white cross on a black background was presented, and participants were instructed to keep their eyes open and let their mind wander. Blood glucose and ketone levels were measured three times throughout the experiment: at baseline, 10 minutes following the bolus, and approximately 60 minutes following the bolus using a Precision Xtra Blood Glucose & Ketone Monitoring System (Abbott Laboratories).

### Structural Magnetic Resonance Imaging (MRI)

Participants were scanned using an ultra-high-field (7T) Siemens Terra MRI Scanner (Siemens Healthineers, Erlangen, Germany) with a 32-channel head coil built in-house at the Massachusetts General Hospital Athinoula A. Martinos Center for Biomedical Imaging. A multi-echo magnetization prepared rapid gradient echo (MEMPRAGE) sequence was used to acquire T1-weighted structural images using a 1 mm isotropic voxel size and four echoes (TE1 = 1.61 ms, TE2 = 3.47 ms, TE3 = 5.33 ms, TE4 = 7.19 ms, TR = 2,530 ms, flip angle = 7°, R = 2 acceleration in the primary phase encoding direction with 32 reference lines), and online GRAPPA image reconstruction, leading to a total volume acquisition time of 6 minutes and 3 seconds.

### ^1^H Magnetic Resonance Imaging (MRS)

^1^H MRS data were collected using a single-voxel stimulated echo acquisition mode (STEAM) sequence with short echo time (TE) = 5.00 ms, repetition time (TR) = 4500 ms, mixing time (TM) = 75 ms, water suppression bandwidth = 132 Hz, averages = 80, voxel size = 20 *×*20 *×*20 mm^3^, acquisition bandwidth = 4000 Hz, vector size = 2048 points, RF pulse duration = 3200 ms, and with 3D outer volume suppression interleaved with variable power and optimized relaxation (VAPOR) water suppression. As our cohort was sampled from across the lifespan, and T_2_ shortens with age in certain metabolites [32], scanning was implemented using a short echo time to avoid underestimating concentration levels. Unsuppressed water acquisition was obtained with similar parameters using four averages. Using the T1-weighted structural image collected prior to MRS acquisition, the voxel was placed in the posterior cortex, encompassing parts of the median posterior cingulate cortex and precuneus, as this region is known to be metabolically active even into old age and well connected to other brain regions [33, 21]. Frequency and eddy current corrections were performed prior to data processing. Concentration levels were estimated using LCModel [43]. Fitted spectra were individually visually inspected for quality and excluded if the spectra contained large lipid peaks or excessive noise. Following the recommendations in [26], the distributions of Cramér-Rao lower bound percent standard deviations (CRLB %SD) were plotted and inspected for each metabolite under each metabolic condition. A CRLB %SD cutoff of 40% was chosen to both ensure reliable quantifications and avoid the rejection of potentially interesting results from metabolites that naturally occur in low concentrations. More details on the CRLB %SD of quantified metabolites can be found in Table 3. Metabolites were analyzed in absolute concentration levels (relative to water) rather than normalized to total creatine (creatine + phosphocreatine, Cr + PCr) due to the changes in the latter observed following the administration of the ketone monoester bolus. Using the T1-weighted structural image, segmentation of the MRS voxel to obtain tissue fraction of gray matter (GM), white matter (WM), and cerebrospinal fluid (CSF) was performed using Gannet (version 3.3.2) [10] and SPM 12 [2]. These tissue fractions were used to perform CSF-correction prior to statistical analyses. GABA concentrations were corrected for CSF based on the GM and WM fractions of the group average voxel fractions, as described in [17],

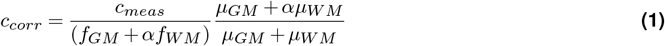

with *α* taken to be 0.5. More details on the group average GM, WM, and CSF fractions can be found in Table 4. The LCModel basis set used for quantification, described in [3], included a D-*β*HB resonance and was used in prior works with ultra-high-field MRS [20]. Averaged spectra macromolecules using metabolite nulling inversion-recovery experiments (acquisition parameters: TR = 2 s and inversion time, TI = 675 ms) were also included in the 7T LCModel basis set as described in [3]. Acquisition parameters have been reported following the guidelines described in [30].

**Table 3.** Distributions of CRLB %SD for each metabolite per condition. If a metabolite was quantified to have a concentration of 0 (meaning a CRLB %SD of 999%) it was removed prior to the calculation of mean and standard deviation in the table above. Value in parentheses indicates the number of quantifications that met the 40% SD threshold as described in Methods.

| Metabolite | Distributions of CRLB %SD |  |  |  |
| --- | --- | --- | --- | --- |
|  | pre-BHB<br>(N = 59) | post-BHB<br>(N = 59) | pre-GLC<br>(N = 56) | post-GLC<br>(N = 57) |
| Asc | 48.1 ± 65.7% (36) | 19.3 ± 21.1% (55) | 53.4 ± 82.4% (38) | 37.1 ± 60.8% (44) |
| Asp | 14.7 ± 7.8% (58) | 18.2 ± 9.3% (58) | 17.6 ± 16.0% (51) | 19.0 ± 30.4% (55) |
| Cr | 6.0 ± 2.7% (59) | 8.3 ± 6.4% (58) | 5.9 ± 2.5% (56) | 6.4 ± 3.5% (57) |
| Cr+PCr | 1.4 ± 0.5% (59) | 1.6 ± 0.7% (59) | 1.4 ± 0.6% (56) | 1.5 ± 0.6% (57) |
| D-βHB | 49.7 ± 67.4% (29) | 6.7 ± 2.8% (59) | 45.5 ± 47.4% (27) | 56.2 ± 120.8% (28) |
| GABA | 12.7 ± 11.9% (57) | 46.3 ± 98.9% (42) | 13.0 ± 13.7% (54) | 14.1 ± 9.7% (55) |
| Glu | 2.1 ± 0.4% (59) | 2.5 ± 1.0% (59) | 2.2 ± 0.5% (56) | 2.2 ± 0.7% (57) |
| Glu+Gln | 2.1 ± 0.7 (59) | 2.5 ± 1.1 (59) | 2.1 ± 0.6 (56) | 2.1 ± 0.8 (57) |
| Gln | 5.2 ± 3.0% (59) | 6.2 ± 2.8% (59) | 4.9 ± 1.9% (56) | 5.3 ± 2.4% (57) |
| GSH | 8.1 ± 2.9% (59) | 8.7 ± 3.6% (59) | 8.2 ± 3.4% (56) | 10.0 ± 8.2% (56) |
| Glc | 110.4 ± 147.7% (10) | 88.8 ± 96.6% (17) | 102.3 ± 109.3% (9) | 41.9 ± 47.7% (35) |
| Glc+Tau | 10.0 ± 3.1% (59) | 12.2 ± 4.5% (59) | 10.1 ± 3.8% (56) | 8.9 ± 3.7% (57) |
| GPC | 7.3 ± 2.2% (59) | 4.9 ± 2.2% (59) | 7.3 ± 2.4% (56) | 7.7 ± 2.3% (57) |
| Ins | 2.6 ± 0.7% (59) | 2.9 ± 1.0% (59) | 2.5 ± 0.8% (56) | 2.7 ± 1.0% (57) |
| Lac | 12.1 ± 7.0% (53) | 17.8 ± 20.5% (47) | 20.9 ± 43.7% (48) | 12.7 ± 8.2% (54) |
| Mac | 2.0 ± 0.6% (59) | 2.2 ± 0.8% (59) | 2.0 ± 0.7% (56) | 2.1 ± 0.5% (57) |
| NAA | 1.6 ± 0.6% (59) | 1.8 ± 0.6% (59) | 1.6 ± 0.5% (56) | 1.6 ± 0.5% (57) |
| NAAG | 9.5 ± 4.2% (58) | 11.2 ± 4.4% (58) | 10.5 ± 7.8% (55) | 10.8 ± 7.4% (55) |
| NAA+NAAG | 1.2 ± 0.4% (59) | 1.3 ± 0.5% (59) | 1.2 ± 0.5% (56) | 1.2 ± 0.5% (57) |
| PCho | 47.6 ± 123.5% (49) | 110.2 ± 153.3% (5) | 33.4 ± 32.2% (45) | 54.4 ± 134.2% (41) |
| PCho+GPC | 4.2 ± 1.4% (59) | 4.5 ± 1.6% (59) | 4.1 ± 1.2% (56) | 4.4 ± 1.7% (57) |
| PCr | 5.6 ± 2.2% (59) | 6.2 ± 2.6% (59) | 5.7 ± 2.1% (56) | 6.0 ± 2.0% (57) |
| PE | 8.9 ± 3.8% (59) | 13.3 ± 12.1% (56) | 9.1 ± 4.2% (56) | 9.7 ± 4.2% (57) |
| sIns | 21.1 ± 13.2% (52) | 27.9 ± 28.7% (45) | 23.2 ± 27.2% (51) | 23.5 ± 18.2% (46) |
| Tau | 9.2 ± 3.0% (59) | 12.6 ± 7.5% (58) | 9.0 ± 3.1% (56) | 9.1 ± 2.7% (57) |

**Table 4.**
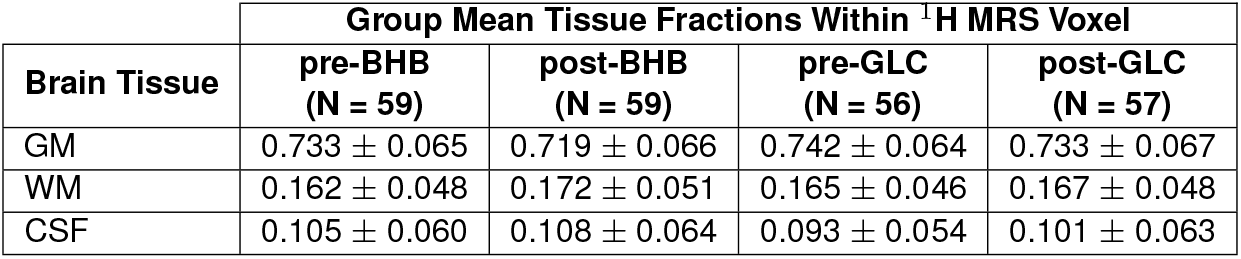
Group means of fractions of gray matter (GM), white matter (WM), and cerebrospinal fluid (CSF) calculated following segmentations of the ^1^**H MRS voxels**. Values are given as mean *±* standard deviation.

### Functional Magnetic Resonance Imaging (fMRI)

Functional MRI data were acquired using a protocol that included whole-brain BOLD (echoplanar imaging, or EPI) images and field map images. The BOLD images were captured using a protocol optimized using a dynamic phantom for detecting resting-state networks with a dynamic phantom (BrainDancer; ALA Scientific Instruments) [27], which included a simultaneous multi-slice (SMS) slice acceleration factor of 5, R=2 acceleration in the primary phase encoding direction (48 reference lines), and online generalized autocalibrating partially parallel acquisition (GRAPPA) image reconstruction. Key acquisition parameters for BOLD imaging were a TR = 802 ms, TE = 20 ms, flip angle = 33^*?*^, voxel size = 2 *×*2 *×*1.5 mm, and a total of 85 slices with 740 measurements (10 minutes) for each pre- and post-bolus interval. The T1-weighted structural images were biasfield corrected, skull-stripped, and normalized to Montreal Neurological Institute (MNI) templates. BOLD contrast functional images obtained from each participant were slice-time corrected and realigned to account for head movement. The BOLD fMRI images were also adjusted for geometric distortions caused by magnetic field inhomogeneity by coregistering the BOLD reference to the intensity-inverted T1-reference [59]. The images were then coregistered with the anatomical images and normalized to MNI space. Mean signals from WM and CSF voxels were regressed out from all time-series to reduce physiological confounds, and six motion regressors were included to minimize motion-related artifacts. The data was spatially smoothed using a kernel with a full width at half maximum (FWHM) of 5 mm.

### Brain Network Instability

Brain network instability is a measure used to describe the persistence of brain networks over time which has been utilized and described in detail in prior work [37, 56]. It can be considered a measure of dynamic functional connectivity [22]. To compute brain network instability, the pre-processed rsfMRI data underwent additional processing steps. First, the clean voxel-space BOLD time-series were band-pass filtered between 0.04 and 0.1 Hz. Next, the filtered time-series were parceled according to the Seitzman functional region of interest (ROI) atlas [50], which included the posterior cingulate cortex (PCC). The parceled time-series were then segmented into non-overlapping 24-second time windows. For each time window, all-to-all signed correlations were calculated using the Ledoit-Wolf covariance estimator. Difference matrices were computed between consecutive time windows and then reduced to a single scalar for each time window pair using an L2 norm. Finally, averaging across all snapshot pairs produced a single scalar value per acquisition, representing whole brain network instability.

### Statistical Analyses

To assess whether the ketone monoester bolus had a significant effect on metabolite concentrations, we first used Shapiro-Wilk tests to evaluate the normality of the distributions for each metabolite, both before and after bolus consumption. Since the majority of these distributions (16 out of 26) deviated significantly from normality, we employed non-parametric Wilcoxon signed-rank tests to compare the metabolite concentrations before and after consumption of the ketone monoester bolus (Fig. 3, Table 2). To investigate the relationship between changes in brain network instability (ΔInstability) and changes in metabolite concentrations (ΔConcentration) following the consumption of glucose and ketone monoester boluses, Spearman correlations were employed. This analysis focused specifically on those metabolites that exhibited statistically significant changes following the ketone monoester bolus administration, as seen in Fig. 3, allowing us to concentrate on the most relevant metabolic shifts and their potential impact on brain network dynamics.

## Results

### Changes in Blood and Brain Concentrations of D-*β*HB and Glucose Following Bolus Consumption are Coupled

Following consumption of the ketone monoester bolus, there is a marked increase in both brain D-*β*HB concentration (Fig. 2A) and blood concentration (Fig. 2B). The same is true for brain and blood glucose (Glc) concentration (Figs. 2D and 2E) following consumption of the glucose bolus. The relationship between brain concentrations and blood concentrations are coupled for both D-*β*HB and Glc following consumption of their respective boluses (Figs. 2C and 2F). Interestingly, we found a significant positive correlation between the change in brain Glc and the change in brain D-*β*HB concentrations following the ketone bolus (N = 55, r = 0.294, p = 0.029), but no significant correlation between the changes in blood Glc and blood D-*β*HB levels following the same bolus (N = 55, r = *−*0.104, p = 0.450).

**Figure 1.**
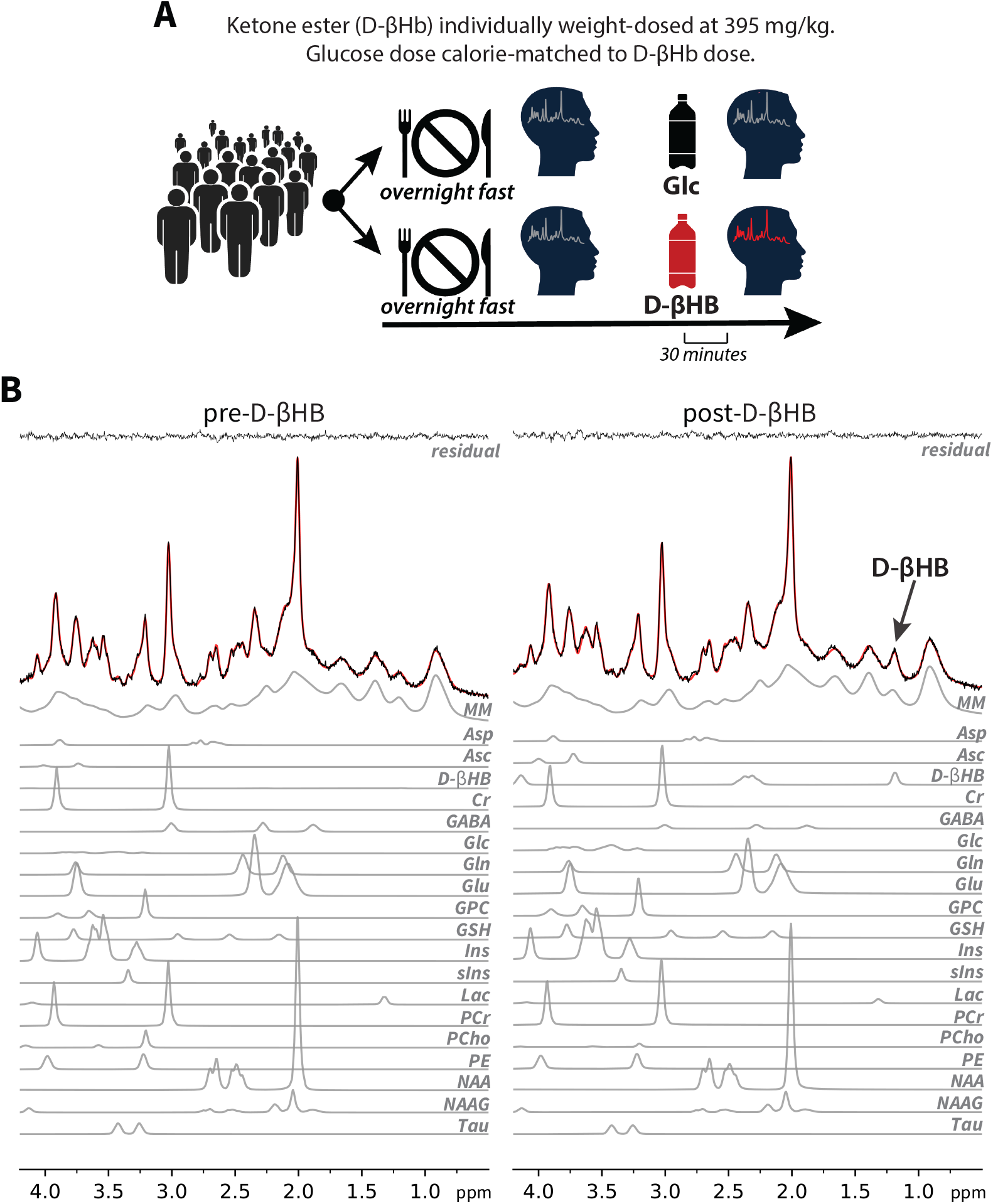
Schematic of experimental design and example spectra. (A) *Design of within-subject, time-locked targeted metabolic* ^1^*H MR Spectroscopy (MRS) experiment*. To quantify the neurochemical effects of oral ketone administration, N = 63 healthy participants from across the lifespan underwent four ^1^H MRS and four rsfMRI scans separated over two days. Following an overnight fast, participants were scanned at baseline and again 30 minutes after consuming a weight-dosed (395 mg/kg) ketone monoester or calorie-matched glucose bolus. The scans were then repeated using the opposite (ketotic or glycolytic) condition on the second day. (B) *Sample* ^1^*H MRS spectra pre- and post-ketone monoester bolus from one participant, fitted using LCModel*. The increase in D-*β*-hydroxybutyrate at 1.2 ppm is clearly visible.

**Figure 2.**
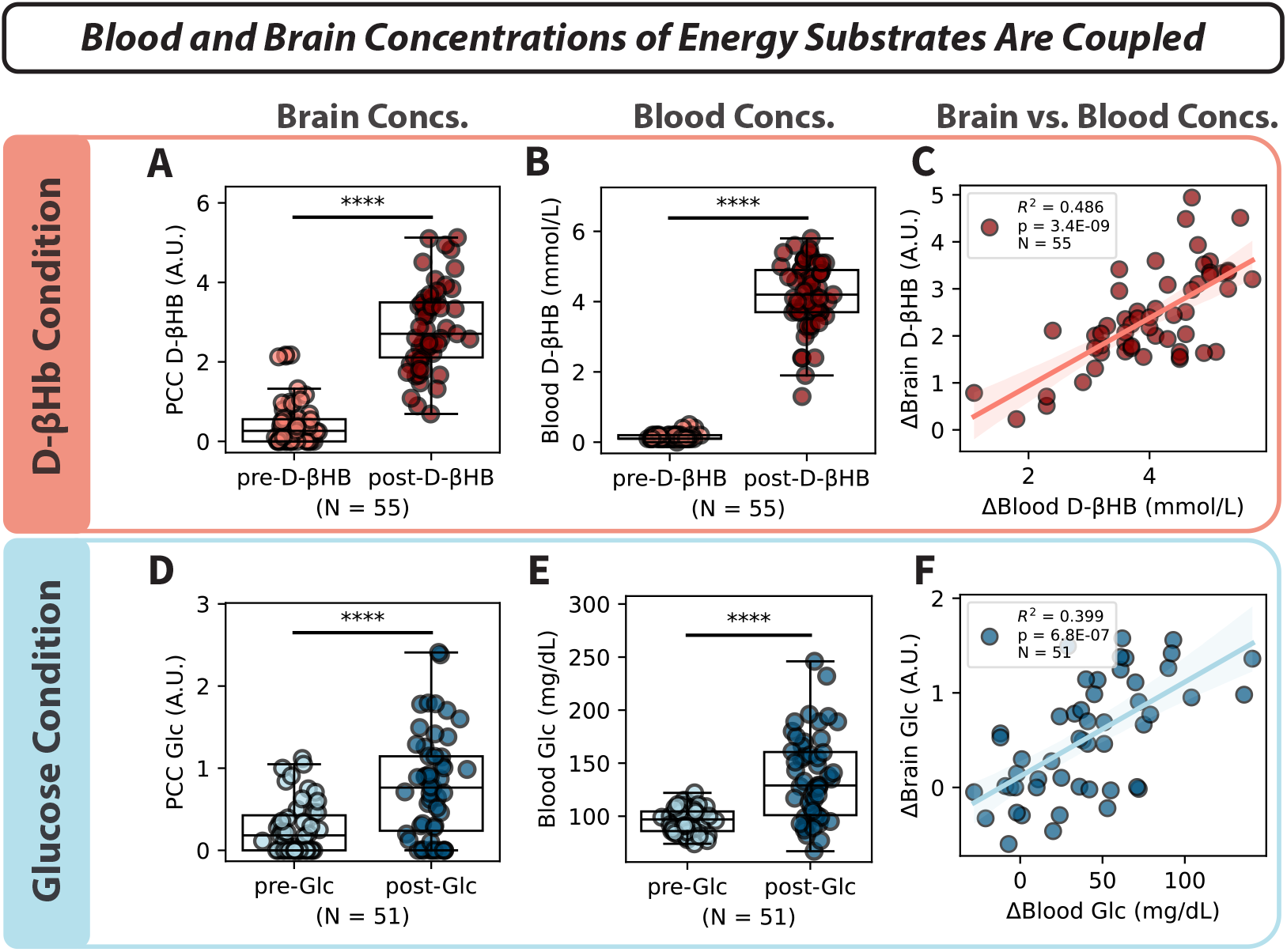
Relationship between metabolite concentrations in the blood and the brain following respective boluses, as well as the effect of creatine changes on brain network stability following the ketone bolus. (A) The concentration of D-*β*HB in the posterior cingulate cortex (PCC) increases markedly following the oral consumption of an exogenous ketone monoester bolus. (B) Following the same bolus, blood concentration of D-*β*HB increase to levels indicating ketosis. (C) The changes in brain and blood D-*β*HB concentrations following the ketone monoester bolus are tightly correlated. (D)-(F) Same as (A)-(C), except now showing the changes in brain and blood glucose (Glc) concentrations following a calorically-matched oral glucose bolus administered on a separate day.

**Figure 3.**
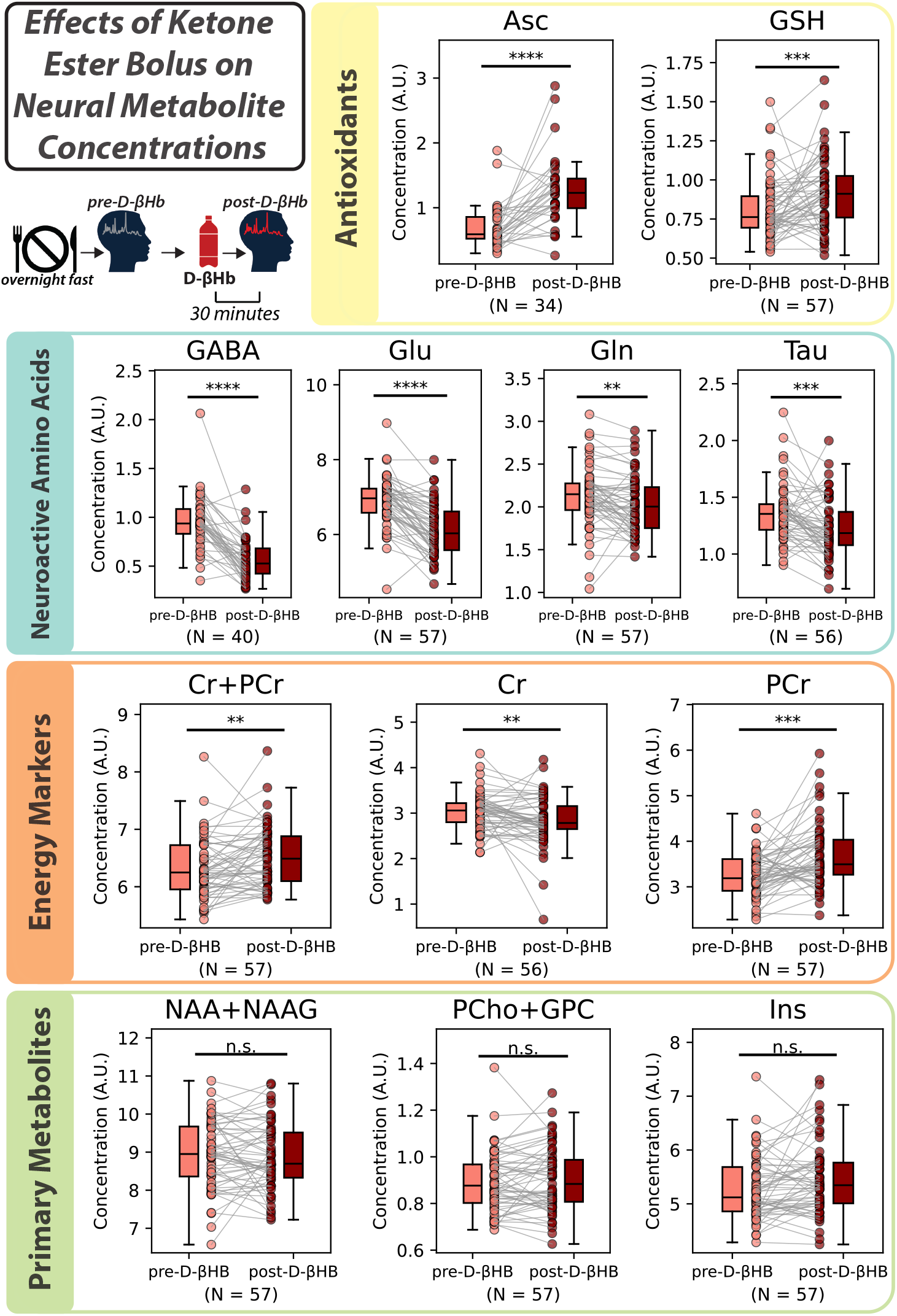
Widespread neural metabolite changes following ketone monoester bolus. Following oral ketone monoester supplementation, we observed significant increases in the antioxidants ascorbate (Asc) and glutathione (GSH). There were notable decreases in the inhibitory neurotransmitter *γ*-aminobutyric acid (GABA) and the excitatory neurotransmitter glutamate (Glu), and slight decreases in glutamine (Gln) and taurine (Tau). Additionally, we observed an increase in total creatine (Cr+PCr), a slight decrease in creatine (Cr), and a significant increase in phosphocreatine (PCr). The basic metabolites N-acetylaspartate+N-acetylaspartylglutamate (NAA+NAAG, or total NAA), phosphocholine+glycerophosphocholine (PCho+GPC, or total cholines), and myo-inositol (Ins) remained unchanged. These results were unique to the ketotic condition and were not observed following glucose supplementation.

### Widespread Changes in Metabolite Concentrations Following Ketones, but not Glucose

Following the ketone monoester bolus, we observed significant increases in the antioxidants ascorbate (Asc) and glutathione (GSH). There were notable decreases in the inhibitory neurotransmitter *γ*-aminobutyric acid (GABA) and the excitatory neurotransmitter glutamate (Glu), and slight decreases in glutamine (Gln) and taurine (Tau). Additionally, we observed a slight increase in total creatine (Cr+PCr), a slight decrease in creatine (Cr), and a significant increase in phosphocreatine (PCr). The basic metabolites N-acetylaspartate+N-acetylaspartylglutamate (NAA+NAAG, or total NAA), phosphocholine+glycerophosphocholine (PCho+GPC, or total cholines), and myo-inositol (Ins) remained unchanged. In contrast, following the glucose bolus, the only significant changes were large increases in Glc and Glc+Tau, and a slight increase in Tau. The changes following the ketone monoester bolus are illustrated in Figure 3, with descriptive statistics for all results outlined in Table 2.

### Higher Fasting Brain Glucose Levels Associated with Increased Neuroinflammation Markers

As the time between bolus administration and MRS measurement was relatively short, we did not expect to observe changes in Ins levels following bolus consumption in our study, as the shortest timescale in published literature over which brain Ins changed following an intervention was four hours, measured in response to lipopolysaccharide administration in an Alzheimer’s disease mouse model [61]. Instead, we analyzed the relationship between fasting brain glucose levels and fasting brain Ins levels, correcting for age, as both fasting glucose (Glc) and Ins increased with age within our cohort (Fig. S1). Our analysis showed a significant positive correlation between fasting Glc and Ins levels (N = 49, r = 0.417, p = 0.003) and an even stronger positive correlation between fasting Glc+Tau and Ins levels (N = 61, r = 0.465, p = 1.579*E −* 04).

### Increase in Brain Bioenergetics Following Ketones Leads to Stabilization of Brain Networks

In the ketotic condition, we observed a negative correlation between change in brain network instability (ΔInstability) and change in Cr+PCr concentration (ΔCr+PCr) (*N* = 55, r = *−* 0.280, p = 0.038, Fig. 4B), meaning individuals who showed greater increases in neural Cr+PCr levels following ketone administration also exhibited more pronounced brain network stabilization.

**Figure 4.**
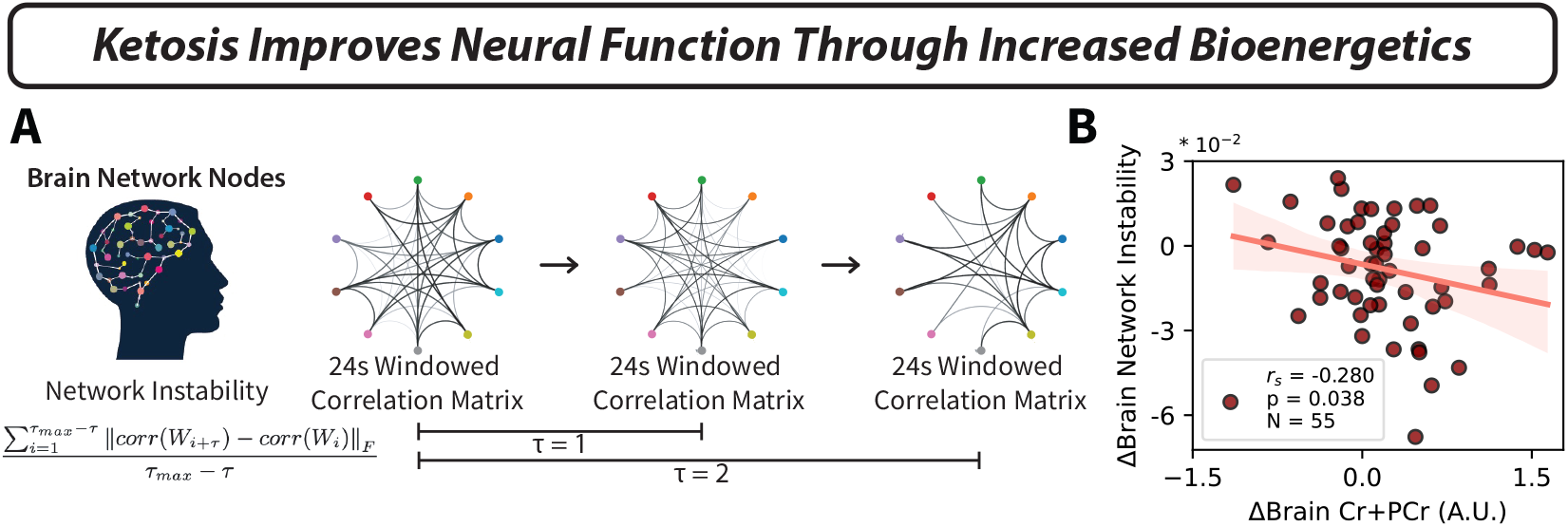
Increase in brain bioenergetics following ketosis leads to an improvement in neural function. (A) *Schematic characterization of brain network instability*. To calculate brain network instability, non-overlapping sliding window correlations are calculated over the entire rsfMRI time-series, with strong correlations defining networks. The stability of the networks is then defined as the degree to which these networks persist over time (in units of *τ* ). (B) *An increase in total neural creatine, as measured using* ^1^*H MR Spectroscopy in the PCC of healthy adults, is associated with a decrease in brain network instability in the same cohort following ketone monoester supplementation*. Both metrics are presented as the difference (Δ) between post-D-*β*HB and pre-D-*β*HB values.

### Changes In Excitatory/Inhibitory Neurotransmitter Ratios Following Ketones

In addition to the overall reductions in GABA and Glutamate observed post-ketone monoester bolus, we observed a significant increase in the Glu/GABA ratio and a decrease in the Glu/Gln ratio (Fig. 5). These changes were specific to the ketone monoester bolus and were not observed following the glucose bolus.

**Figure 5.**
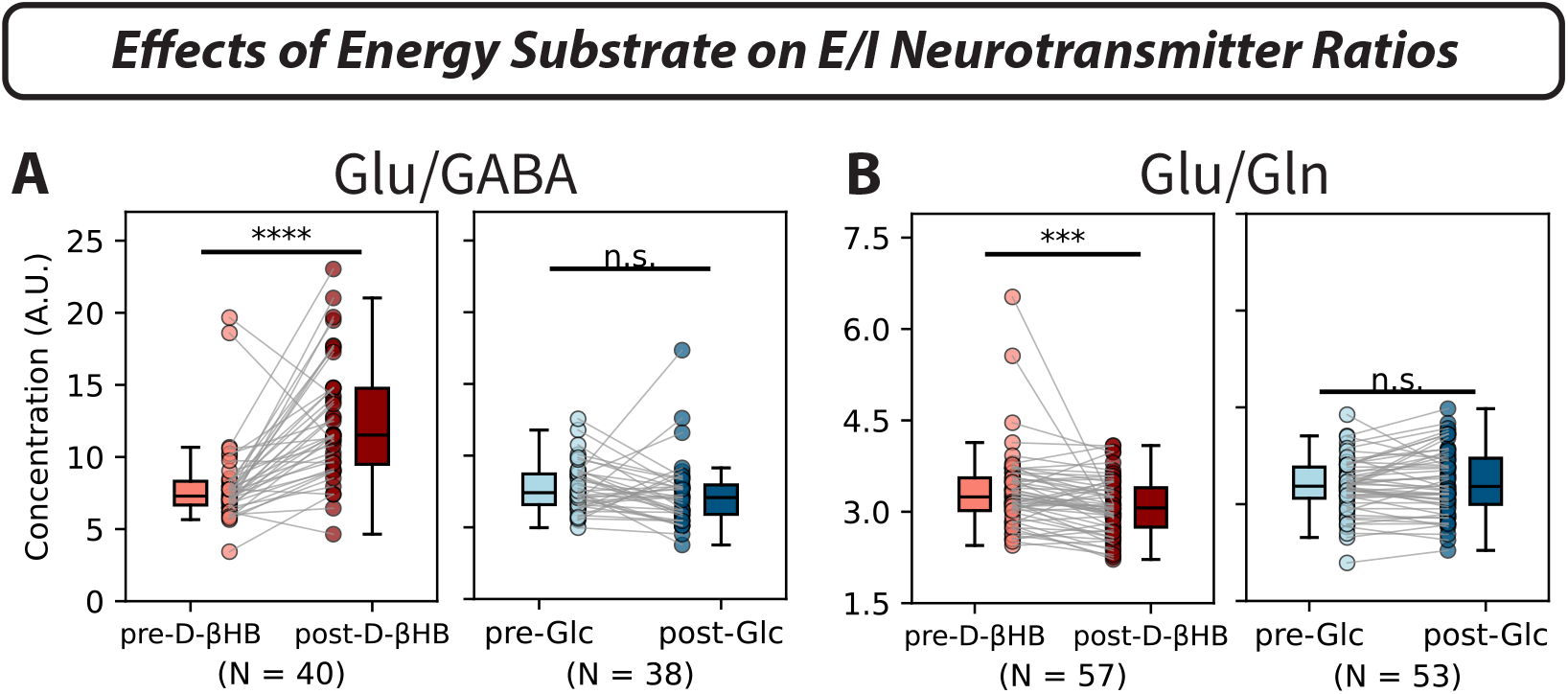
The ratios of excitatory (Glu) to inhibitory (GABA) neurotransmitters and neurotransmitter precursor (Gln) change following the ketone monoester bolus, but remain unchanged following the glucose bolus in the PCC of healthy adults. (A) The ratio of Glu/GABA increased significantly following consumption of an exogenous ketone monoester bolus, but not following a calorie-matched glucose bolus. (B) The ratio of Glu/Gln decreased following the ketone monoester bolus but not following the glucose bolus.

## Discussion

The current study significantly expands upon our previous study by increasing the sample size from 26 to 63 participants, allowing for a more robust examination of the neurochemical effects of ketone and glucose administration across a broad age range and the capacity to relate these neurochemical changes to functional changes. We replicated earlier findings of decreased *γ*-Aminobutyric Acid (GABA) and glutamate (Glu) levels following ketone administration [20], but also identified additional changes in brain metabolites not observed previously (Figs. 3, 5). Specifically, we observed significant increases in the antioxidants glutathione (GSH) and ascorbate (Asc). We observed decreases in glutamine (Gln), taurine (Tau), the GABA/Glu ratio, and the Glu/Gln ratio, and increases in the Glu/GABA and Gln/Glu ratios. We also observed a positive correlation between fasting brain glucose (Glc) levels and fasting brain myo-inositol (Ins) levels. Finally, connecting our ^1^H MRS results to our fMRI data taken from same cohort pointed to an improvement in brain bioenergetics (in the form of an increase in total creatine) as a potential mechanism for brain network stabilization observed following consumption of ketone monoester [37].

The loss of glucose homeostasis in the elderly and its correlation with myoinositol levels supports a relationship with inflammation. Fluctuations in glucose levels can lead to the activation of microglia, increasing oxidative stress and leading to neuroinflammation and potential cognitive dysfunction [60], while high glucose levels have been found to induce neuroinflammation and apoptosis in the brain, particularly in the context of traumatic brain injury (TBI) [63]. Myo-inositol (Ins) is higher in concentration in astrocytes than in neurons [7] and as such is considered to be an astroglial marker and a subsequent marker of neuroinflammation [9]. Ins is elevated in those with type-2 diabetes [14, 49] and those with hyperglycemia due to type 1 diabetes [19], suggesting a link between dysregulated glucose metabolism and neuroinflammation. Additionally, elevated Ins levels have been observed in patients with mild cognitive impairment and Alzheimer’s disease [16, 57], and in veterans with a history of TBI [52].

In our healthy cohort, we observed a significant positive correlation between fasting brain glucose levels and Ins while correcting for age, suggesting that glucose itself may be associated with inflammatory processes and contribute to neuroinflammation even before a person enters a diseased state. Several potential mechanisms could explain this observation. High glucose levels can lead to an increased production of advanced glycation end-products (AGEs), which are known to induce oxidative stress and inflammation [53, 35]. Chronic hyperglycemia can also lead to alterations in brain metabolism, cellular signaling pathways, and the blood brain barrier, contributing to neuroinflammation [31, 42].

Beyond neuroinflammation, the brain in particular is especially sensitive to oxidative stress due to its high metabolic activity and low antioxidant capacity [29, 23], which in turn contributes to neurodegenerative diseases and age-related cognitive decline [11]. We observed increases in the antioxidants ascorbate and glutathione following the ketone monoester bolus. Ascorbate, primarily found in neurons, and glutathione, primarily found in astrocytes [46], both directly scavenge reactive oxygen species (ROS) [18], which are known to induce oxidative stress if the ratio between ROS production and elimination becomes unbalanced [4]. We also observed a small decrease in taurine. However, accurately distinguishing glucose from taurine using ^1^*H* MRS can be challenging due to their spectral proximity, and we did not observe a change in Glc+Tau following the ketone bolus 2. The separate contributions of Glc and Tau to the combined concentrations can be determined at 7T via editing techniques, and further studies will be needed before a conclusive assignment can be made. Lastly, we observed an increase in total creatine (creatine plus phosphocreatine, or Cr+PCr) following the ketone bolus, which is consistent with its well-established role in energy metabolism. Creatine metabolism is especially significant during periods of rapid increases in neuronal activity, such as those occurring during sensory and cognitive activation, and has been shown to be impaired in bipolar disorder [62]. Furthermore, impaired bioenergetics have been found to be associated with various psychiatric disorders [55]. In line with these findings, creatine supplementation has been reported to improve cognitive performance [15], underscoring its importance in neurological function. It should be noted that this observed increase in total creatine makes the usual method of scaling concentrations to Cr+PCr invalid, and should be accounted for in the field when the use of exogenous ketones or ketogenic diets are included in the experimental design.

In addition to changes in bioenergetics, we observed changes in the concentrations and balances of excitatory and inhibitory neurotransmitters following acute ketosis. The glutamate/GABA-glutamine cycle plays a critical role in maintaining the homeostasis of glutamate and GABA in the brain [5]. Dysfunction in the glutamate-glutamine cycle has been implicated in various neurological disorders, including Alzheimer’s disease [58], epilepsy [41], and bipolar disorder [54]. Our findings of reduced GABA and glutamate following ketone monoester bolus administration correspond to those previously reported [20]. However, our increased sample size enabled us to detect a significant reduction in glutamine levels, an effect not observed previously.

The simultaneous reduction in glutamate, GABA and glutamine indicates an overall decrease in the total neurotransmitter pool. This reduction could be explained by a slowdown in neuronal glucose metabolism caused by ketones. Neurotransmitter cycling and glycolysis have been shown to follow a 1:1 flux ratio, likely because of their coupling through cytosolic redox balance [28, 47, 48]. Glucose, the main energy source for neurons, undergoes glycolysis. Downstream to glycolysis, pyruvate is produced, which is converted into acetyl-CoA, a key substrate for the TCA cycle. Redox equivalents generated by glycolysis are transferred to mitochondria by the malate aspartate shuttle, where it provides energy for mitochondrial glutamate synthesis from glutamine and transports it back to the cytosol. Once in the cytosol, it is packaged into vesicles and released from nerve terminals during depolarization. GABA is synthesized from glutamine via a similar pathway [48]. While glucose is metabolized into acetyl-CoA through glycolysis, D-*β*HB is also converted to acetyl-CoA, but via a pathway that is different from glycolysis and thus does not affect cytosolic redox conditions. Consequently, ketones do not contribute to driving the neurotransmitter flux. When both glucose and ketones are available in substantial amounts, ketones produce sufficient acetyl-CoA to compete with glucose for the limited capacity of TCA cycle enzymes [40, 24]. This competition may result in a backlog in glucose-derived pathways that produce acetyl-CoA, including glycolysis, which in turn slows the synthesis of glutamate and GABA via the malate aspartate shuttle.

The increase in the antioxidants ascorbate and glutathione following ketone consumption may be explained by the interplay between glycolysis and the parallel pentose phosphate pathway (PPP). Inhibition of glycolysis can lead to a compensatory increase in flux through the PPP due to the accumulation of glucose-6-phosphate (G6P). G6P is then shunted into the oxidative phase of the PPP, producing NADPH, which in turn can reduce oxidized glutathione (GSSG) back to glutathione via glutathione reductase [13]. While ascorbate is not directly coupled with the PPP and NADPH, dehydroascorbate (DHA) is reduced into ascorbate by GSH, which typically results in an inverse relationship between ascorbate levels and GSH/GSSG ratios [6]. However, it can be hypothesized that an overall increase in GSH availability may lead to enhanced oxidation of DHA into ascorbate, increasing the overall amounts of these two closely linked antioxidants.

With this context in mind, our results demonstrate the neuroprotective utility of ketones as an alternative fuel source in three ways. First, the utilization of ketones instead of glucose avoids an increase in glucose levels in both the brain and in the periphery (Table 2, Fig. 2). This avoids potential neuroinflammation caused by fluctuations in and high levels of glucose, especially over long periods of time, while still providing the brain with the energy needed to function. Second, ketones increase the levels of antioxidants in the brain, which may prevent oxidative damage that occurs as a natural byproduct of metabolism and aging, and may be helpful in treating or preventing neurodegenerative and neuropsychiatric disorders associated with oxidative stress or mitochondrial dysfunction. Finally, ketones enhance brain function partly by increasing creatine levels, which improves brain bioenergetics and subsequently stabilize brain networks as measured using fMRI. The combination of metabolic and functional neuroimaging data in our study provides a comprehensive view of how ketosis affects brain metabolite concentrations and subsequently functional network dynamics, providing insights necessary for developing novel treatment strategies for a variety of neurodegenerative and psychiatric disorders.

## Acknowledgements

We would like to thank Dr. Dinesh Deelchand from the Center for Magnetic Resonance Research at the University of Minnesota for providing us with the 7T LCModel basis set used in our analysis, and Dr. Akila Weerasekera for help with the basis set. The research was funded by the W. M. Keck Foundation (to L.R.M.-P.) and the NSF Brain Research through Advancing Innovative Neurotechnologies (BRAIN) Initiative NCS-FR 1926 781 (to L.R.M.-P.).

## Supplementary Information

**Figure S1.**
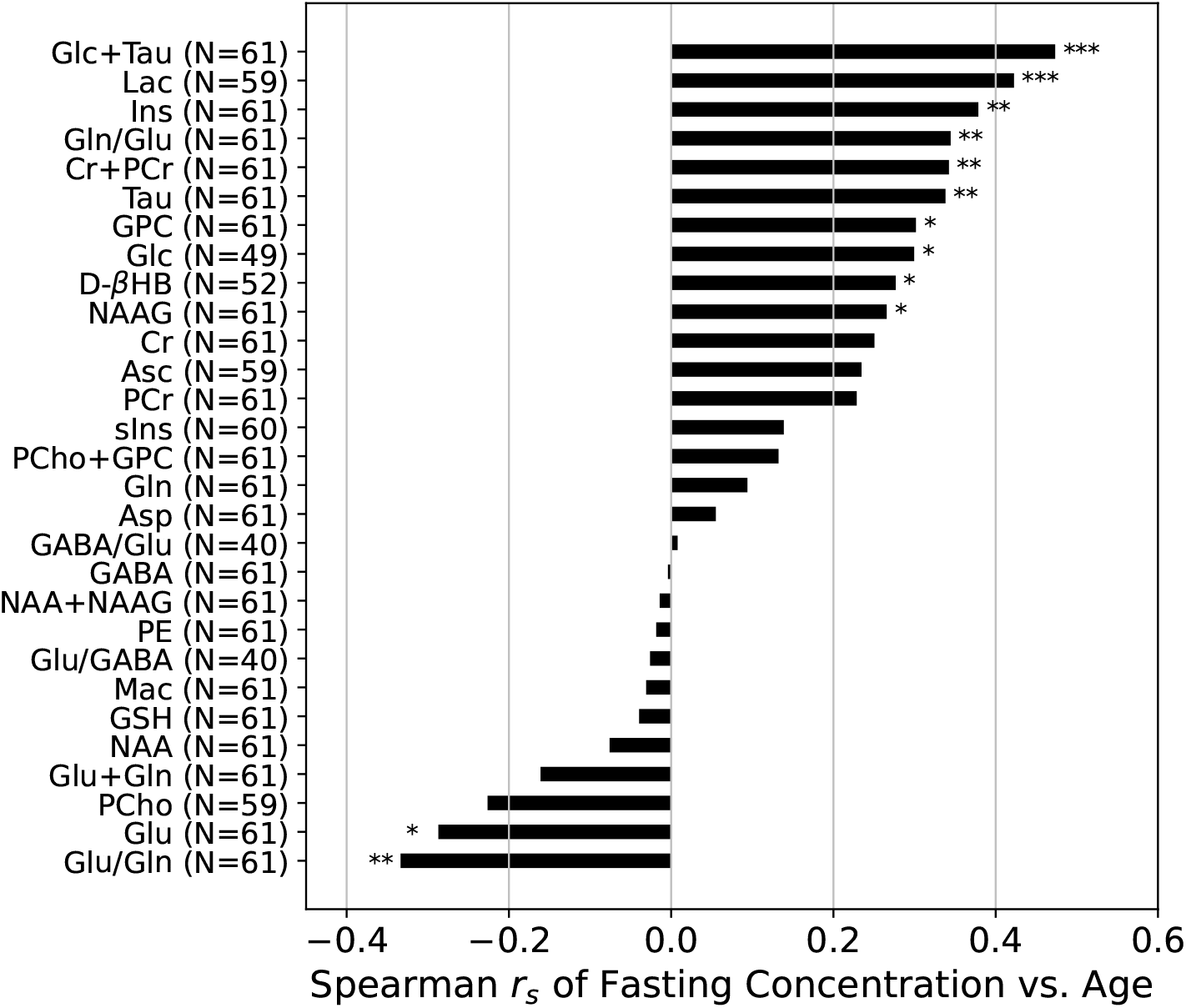
Plots of Spearman correlation coefficients (*r*_*s*_-values) between pre-bolus metabolite concentrations and age. Only spectra with CRLB %SD < 300% were included in the analysis. Fasting concentration values were obtained by averaging pre-D-*β*HB and pre-Glc values when both were available, otherwise, either pre-D-*β*HB or pre-Glc was used as the value. Asterisks indicate the p-values of the correlations.

## Bibliography

1. Botond Antal, Liam P McMahon, Syed Fahad Sultan, Andrew Lithen, Deborah J Wexler, Bradford Dickerson, Eva-Maria Ratai, and Lilianne R Mujica-Parodi. Type 2 diabetes mellitus accelerates brain aging and cognitive decline: Complementary findings from uk biobank and meta-analyses. Elife, 11:e73138, 2022.

2. John Ashburner and Karl J Friston. Unified segmentation. neuroimage, 26(3):839–851, 2005.

3. Nazem Atassi, Maosheng Xu, Christina Triantafyllou, Boris Keil, Robert Lawson, Paul Cernasov, Elena Ratti, Christopher J Long, Sabrina Paganoni, Alyssa Murphy, et al. Ultra high-field (7tesla) magnetic resonance spectroscopy in amyotrophic lateral sclerosis. PLoS One, 12(5):e0177680, 2017.

4. Richard L Auten and Jonathan M Davis. Oxygen toxicity and reactive oxygen species: the devil is in the details. Pediatric research, 66(2):121–127, 2009.

5. Lasse K Bak, Arne Schousboe, and Helle S Waagepetersen. The glutamate/gaba-glutamine cycle: aspects of transport, neurotransmitter homeostasis and ammonia transfer. Journal of neurochemistry, 98(3):641–653, 2006.

6. Gábor Bánhegyi, László Braun, Miklós Csala, Ferenc Puskás, and József Mandl. Ascorbate metabolism and its regulation in animals. Free Radical Biology and Medicine, 23(5):793–803, 1997.

7. A Brand, C Richter-Landsberg, and D Leibfritz. Multinuclear nmr studies on the energy metabolism of glial and neuronal cells. Developmental neuroscience, 15(3-5):289–298, 1993.

8. Stephen C Cunnane, Eugenia Trushina, Cecilie Morland, Alessandro Prigione, Gemma Casadesus, Zane B Andrews, M Flint Beal, Linda H Bergersen, Roberta D Brinton, Suzanne de la Monte, et al. Brain energy rescue: an emerging therapeutic concept for neurodegenerative disorders of ageing. Nature reviews Drug discovery, 19(9):609–633, 2020.

9. Danielle E Eagan, Mitzi M Gonzales, Takashi Tarumi, Hirofumi Tanaka, Sandra Stautberg, and Andreana P Haley. Elevated serum c-reactive protein relates to increased cerebral myoinositol levels in middle-aged adults. Cardiovascular psychiatry and neurology, 2012(1):120540, 2012.

10. Richard AE Edden, Nicolaas AJ Puts, Ashley D Harris, Peter B Barker, and C John Evans. Gannet: A batch-processing tool for the quantitative analysis of gamma-aminobutyric acid–edited mr spectroscopy spectra. Journal of magnetic resonance imaging, 40(6):1445–1452, 2014.

11. Ferdinando Franzoni, Giorgia Scarfò, Sara Guidotti, Jonathan Fusi, Muzaffar Asomov, and Carlo Pruneti. Oxidative stress and cognitive decline: the neuroprotective role of natural antioxidants. Frontiers in neuroscience, 15:729757, 2021.

12. John M Freeman, Eileen PG Vining, Diana J Pillas, Paula L Pyzik, Jane C Casey, LCSW; Kelly, and Millicent T. The efficacy of the ketogenic diet—1998: a prospective evaluation of intervention in 150 children. Pediatrics, 102(6):1358–1363, 1998.

13. Tongxin Ge, Jiawen Yang, Shihui Zhou, Yuchen Wang, Yakui Li, and Xuemei Tong. The role of the pentose phosphate pathway in diabetes and cancer. Frontiers in Endocrinology, 11:365, 2020.

14. A Geissler, R Fründ, J Schölmerich, S Feuerbach, and B Zietz. Alterations of cerebral metabolism in patients with diabetes mellitus studied by proton magnetic resonance spectroscopy. Experimental and clinical endocrinology & diabetes, 111(07): 421–427, 2003.

15. Ali Gordji-Nejad, Andreas Matusch, Sophie Kleedörfer, Harshal Jayeshkumar Patel, Alexander Drzezga, David Elmenhorst, Ferdinand Binkofski, and Andreas Bauer. Single dose creatine improves cognitive performance and induces changes in cerebral high energy phosphates during sleep deprivation. Scientific reports, 14(1):4937, 2024.

16. H Randall Griffith, Ozioma C Okonkwo, Jan A den Hollander, Katherine Belue, Jacqueline Copeland, Lindy E Harrell, John C Brockington, David G Clark, and Daniel C Marson. Brain metabolic correlates of decision making in amnestic mild cognitive impairment. Aging, Neuropsychology, and Cognition, 17(4):492–504, 2010.

17. Ashley D Harris, Nicolaas AJ Puts, and Richard AE Edden. Tissue correction for gaba-edited mrs: Considerations of voxel composition, tissue segmentation, and tissue relaxations. Journal of Magnetic Resonance Imaging, 42(5):1431–1440, 2015.

18. Fiona E Harrison and James M May. Vitamin c function in the brain: vital role of the ascorbate transporter svct2. Free Radical Biology and Medicine, 46(6):719–730, 2009.

19. O Heikkilä, N Lundbom, M Timonen, P-H Groop, S Heikkinen, and S Mäkimattila. Hyperglycaemia is associated with changes in the regional concentrations of glucose and myo-inositol within the brain. Diabetologia, 52:534–540, 2009.

20. Antoine Hone-Blanchet, Botond Antal, Liam McMahon, Andrew Lithen, Nathan A Smith, Steven Stufflebeam, Yi-Fen Yen, Alexander Lin, Bruce G Jenkins, Lilianne R Mujica-Parodi, et al. Acute administration of ketone beta-hydroxybutyrate downregulates 7t proton magnetic resonance spectroscopy-derived levels of anterior and posterior cingulate gaba and glutamate in healthy adults. Neuropsychopharmacology, 48(5):797–805, 2023.

21. Mingming Huang, Hui Yu, Xi Cai, Yong Zhang, Wei Pu, and Bo Gao. A comparative study of posterior cingulate metabolism in patients with mild cognitive impairment due to parkinson’s disease or alzheimer’s disease. Scientific Reports, 13(1):14241, 2023.

22. R Matthew Hutchison, Thilo Womelsdorf, Elena A Allen, Peter A Bandettini, Vince D Calhoun, Maurizio Corbetta, Stefania Della Penna, Jeff H Duyn, Gary H Glover, Javier Gonzalez-Castillo, et al. Dynamic functional connectivity: promise, issues, and interpretations. Neuroimage, 80:360–378, 2013.

23. Matyas Jelinek, Michal Jurajda, and Kamil Duris. Oxidative stress in the brain: basic concepts and treatment strategies in stroke. Antioxidants, 10(12):1886, 2021.

24. Nicole Jacqueline Jensen, Helena Zander Wodschow, Malin Nilsson, and Jørgen Rungby. Effects of ketone bodies on brain metabolism and function in neurodegenerative diseases. International journal of molecular sciences, 21(22):8767, 2020.

25. Brad Kincaid and Ella Bossy-Wetzel. Forever young: Sirt3 a shield against mitochondrial meltdown, aging, and neurodegeneration. Frontiers in aging neuroscience, 5:58146, 2013.

26. Roland Kreis. The trouble with quality filtering based on relative cramer-rao lower bounds. Magnetic resonance in medicine, 75(1):15–18, 2016.

27. Rajat Kumar, Liang Tan, Alan Kriegstein, Andrew Lithen, Jonathan R Polimeni, Lilianne R Mujica-Parodi, and Helmut H Strey. Ground-truth “resting-state” signal provides data-driven estimation and correction for scanner distortion of fmri time-series dynamics. NeuroImage, 227:117584, 2021.

28. Bernard Lanz, Lijing Xin, Philippe Millet, and Rolf Gruetter. In vivo quantification of neuro-glial metabolism and glial glutamate concentration using 1h-[13c] mrs at 14.1 t. Journal of neurochemistry, 128(1):125–139, 2014.

29. Kyung Hee Lee, Myeounghoon Cha, and Bae Hwan Lee. Neuroprotective effect of antioxidants in the brain. International journal of molecular sciences, 21(19):7152, 2020.

30. Alexander Lin, Ovidiu Andronesi, Wolfgang Bogner, In-Young Choi, Eduardo Coello, Cristina Cudalbu, Christoph Juchem, Graham J Kemp, Roland Kreis, Martin Krššák, et al. Minimum reporting standards for in vivo magnetic resonance spectroscopy (mrsinmrs): experts’ consensus recommendations. NMR in Biomedicine, 34(5):e4484, 2021.

31. Pierre J Magistretti and Igor Allaman. A cellular perspective on brain energy metabolism and functional imaging. Neuron, 86(4):883–901, 2015.

32. Malgorzata Marjańska, Uzay E Emir, Dinesh K Deelchand, and Melissa Terpstra. Faster metabolite 1h transverse relaxation in the elder human brain. PloS one, 8(10):e77572, 2013.

33. Malgorzata Marjańska, J Riley McCarten, James Hodges, Laura S Hemmy, Andrea Grant, Dinesh K Deelchand, and Melissa Terpstra. Region-specific aging of the human brain as evidenced by neurochemical profiles measured noninvasively in the posterior cingulate cortex and the occipital lobe using 1h magnetic resonance spectroscopy at 7 t. Neuroscience, 354: 168–177, 2017.

34. Saroshi Minoshima, Bruno Giordani, Stanley Berent, Kirk A Frey, Norman L Foster, and David E Kuhl. Metabolic reduction in the posterior cingulate cortex in very early alzheimer’s disease. Annals of Neurology: Official Journal of the American Neurological Association and the Child Neurology Society, 42(1):85–94, 1997.

35. Sanne S Mooldijk, Tianqi Lu, Komal Waqas, Jinluan Chen, Meike W Vernooij, Kamran Ikram, M Carola Zillikens, and M Arfan Ikram. Skin autofluorescence, reflecting accumulation of advanced glycation end products, and the risk of dementia in a population-based cohort. Scientific Reports, 14(1):1256, 2024.

36. Gerwyn Morris, Michael Berk, Ken Walder, and Michael Maes. Central pathways causing fatigue in neuro-inflammatory and autoimmune illnesses. BMC medicine, 13:1–23, 2015.

37. Lilianne R Mujica-Parodi, Anar Amgalan, Syed Fahad Sultan, Botond Antal, Xiaofei Sun, Steven Skiena, Andrew Lithen, Noor Adra, Eva-Maria Ratai, Corey Weistuch, et al. Diet modulates brain network stability, a biomarker for brain aging, in young adults. Proceedings of the National Academy of Sciences, 117(11):6170–6177, 2020.

38. Elizabeth G Neal, Hannah Chaffe, Ruby H Schwartz, Margaret S Lawson, Nicole Edwards, Geogianna Fitzsimmons, Andrea Whitney, and J Helen Cross. The ketogenic diet for the treatment of childhood epilepsy: a randomised controlled trial. The Lancet Neurology, 7(6):500–506, 2008.

39. Felicity Ng, Michael Berk, Olivia Dean, and Ashley I Bush. Oxidative stress in psychiatric disorders: evidence base and therapeutic implications. International Journal of Neuropsychopharmacology, 11(6):851–876, 2008.

40. Jullie W Pan, Robin A de Graaf, Kitt F Petersen, Gerald I Shulman, Hoby P Hetherington, and Douglas L Rothman. 2, 4-13c2]-β-hydroxybutyrate metabolism in human brain. Journal of Cerebral Blood Flow & Metabolism, 22(7):890–898, 2002.

41. Ognen AC Petroff, Laura D Errante, Douglas L Rothman, Jung H Kim, and Dennis D Spencer. Glutamate–glutamine cycling in the epileptic human hippocampus. Epilepsia, 43(7):703–710, 2002.

42. Chandan Prasad, Kathleen E Davis, Victorine Imrhan, Shanil Juma, and Parakat Vijayagopal. Advanced glycation end products and risks for chronic diseases: intervening through lifestyle modification. American journal of lifestyle medicine, 13 (4):384–404, 2019.

43. Stephen W Provencher. Estimation of metabolite concentrations from localized in vivo proton nmr spectra. Magnetic resonance in medicine, 30(6):672–679, 1993.

44. Marcus E Raichle, Ann Mary MacLeod, Abraham Z Snyder, William J Powers, Debra A Gusnard, and Gordon L Shulman. A default mode of brain function. Proceedings of the national academy of sciences, 98(2):676–682, 2001.

45. Gislaine T Rezin, Graziela Amboni, Alexandra I Zugno, João Quevedo, and Emilio L Streck. Mitochondrial dysfunction and psychiatric disorders. Neurochemical research, 34:1021–1029, 2009.

46. Margaret E Rice and I Russo-Menna. Differential compartmentalization of brain ascorbate and glutathione between neurons and glia. Neuroscience, 82(4):1213–1223, 1997.

47. Douglas L Rothman, Gerald A Dienel, Kevin L Behar, Fahmeed Hyder, Mauro DiNuzzo, Federico Giove, and Silvia Mangia. Glucose sparing by glycogenolysis (gsg) determines the relationship between brain metabolism and neurotransmission. Journal of Cerebral Blood Flow & Metabolism, 42(5):844–860, 2022.

48. Douglas L Rothman, Kevin L Behar, and Gerald A Dienel. Mechanistic stoichiometric relationship between the rates of neurotransmission and neuronal glucose oxidation: Reevaluation of and alternatives to the pseudo-malate-aspartate shuttle model. Journal of Neurochemistry, 168(5):555–591, 2024.

49. Rajani Santhakumari, Indla Yogananda Reddy, and R Archana. Effect of type 2 diabetes mellitus on brain metabolites by using proton magnetic resonance spectroscopy-a systematic review. International journal of pharma and bio sciences, 5(4): 1118, 2014.

50. Benjamin A Seitzman, Caterina Gratton, Scott Marek, Ryan V Raut, Nico UF Dosenbach, Bradley L Schlaggar, Steven E Petersen, and Deanna J Greene. A set of functionally-defined brain regions with improved representation of the subcortex and cerebellum. Neuroimage, 206:116290, 2020.

51. Shebani Sethi, Diane Wakeham, Terence Ketter, Farnaz Hooshmand, Julia Bjornstad, Blair Richards, Eric Westman, Ronald M Krauss, and Laura Saslow. Ketogenic diet intervention on metabolic and psychiatric health in bipolar and schizophrenia: A pilot trial. Psychiatry research, 335:115866, 2024.

52. Chandni Sheth, Andrew P Prescot, Margaret Legarreta, Perry F Renshaw, Erin McGlade, and Deborah Yurgelun-Todd. Increased myoinositol in the anterior cingulate cortex of veterans with a history of traumatic brain injury: a proton magnetic resonance spectroscopy study. Journal of Neurophysiology, 123(5):1619–1629, 2020.

53. Varun Parkash Singh, Anjana Bali, Nirmal Singh, and Amteshwar Singh Jaggi. Advanced glycation end products and diabetic complications. The Korean journal of physiology & pharmacology: official journal of the Korean Physiological Society and the Korean Society of Pharmacology, 18(1):1, 2014.

54. Márcio Gerhardt Soeiro-de Souza, Anke Henning, Rodrigo Machado-Vieira, Ricardo A Moreno, Bruno F Pastorello, Cláudia da Costa Leite, Homero Vallada, and Maria Concepcion Garcia Otaduy. Anterior cingulate glutamate–glutamine cycle metabolites are altered in euthymic bipolar i disorder. European neuropsychopharmacology, 25(12):2221–2229, 2015.

55. Xiaopeng Song, Xi Chen, Cagri Yuksel, Junliang Yuan, Diego A Pizzagalli, Brent Forester, Dost Öngür, and Fei Du. Bioenergetics and abnormal functional connectivity in psychotic disorders. Molecular psychiatry, 26(6):2483–2492, 2021.

56. Helena van Nieuwenhuizen, Anthony G Chesebro, Claire Polizu, Kieran Clarke, Helmut H Strey, Corey Weistuch, and Lilianne R Mujica-Parodi. Ketosis regulates k+ ion channels, strengthening brain-wide signaling disrupted by age. Imaging Neuroscience, 2:1–14, 2024.

57. Olga Voevodskaya, Konstantinos Poulakis, Pia Sundgren, Danielle Van Westen, Sebastian Palmqvist, Lars-Olof Wahlund, Erik Stomrud, Oskar Hansson, Eric Westman, Swedish BioFINDER Study Group, et al. Brain myoinositol as a potential marker of amyloid-related pathology: A longitudinal study. Neurology, 92(5):e395–e405, 2019.

58. Heather Scott Walton and Peter R Dodd. Glutamate–glutamine cycling in alzheimer’s disease. Neurochemistry international, 50(7-8):1052–1066, 2007.

59. Sijia Wang, Daniel J Peterson, J Christopher Gatenby, Wenbin Li, Thomas J Grabowski, and Tara M Madhyastha. Evaluation of field map and nonlinear registration methods for correction of susceptibility artifacts in diffusion mri. Frontiers in neuroinformatics, 11:17, 2017.

60. Charles Watt, Elizabeth Sanchez-Rangel, and Janice Jin Hwang. Glycemic variability and cns inflammation: Reviewing the connection. Nutrients, 12(12):3906, 2020.

61. Maria Yanez Lopez, Marie-Christine Pardon, Kerstin Baiker, Malcolm Prior, Ding Yuchun, Alessandra Agostini, Li Bai, Dorothee P Auer, and Henryk M Faas. Myoinositol cest signal in animals with increased iba-1 levels in response to an inflammatory challenge—preliminary findings. PLoS One, 14(2):e0212002, 2019.

62. C Yuksel, F Du, C Ravichandran, JR Goldbach, T Thida, P Lin, B Dora, J Gelda, L O’connor, S Sehovic, et al. Abnormal high-energy phosphate molecule metabolism during regional brain activation in patients with bipolar disorder. Molecular psychiatry, 20(9):1079–1084, 2015.

63. Wenqian Zhang, Jun Hong, Wencheng Zheng, Aijun Liu, and Ying Yang. High glucose exacerbates neuroinflammation and apoptosis at the intermediate stage after post-traumatic brain injury. Aging (Albany NY), 13(12):16088, 2021.

